# Chamber Implant for Chronic Optical Recordings from the Cerebral Cortex of Marmosets

**DOI:** 10.1101/2025.07.27.667056

**Authors:** Justine Marguin, Luc Renaud, Xavier Degiovanni, Alberto Lombardini, Rana Banai-Tizkar, Sebastien Roux, Manon Clémenceau, Christophe Melon, Frederic Chavane, Ivo Vanzetta

**Affiliations:** Aix Marseille Univ., CNRS, Institut de Neurosciences de la Timone (INT), UMR 7289, Marseille, France

**Keywords:** Marmoset, Optical imaging, Two-photon microscopy, Chronic recording, Cranial window, Neurotechniques, Genetically encoded fluorescent calcium, voltage sensors

## Abstract

We describe a novel imaging chamber for chronic optical recordings from the marmoset cerebral cortex, together with the surgery needed for its implantation and that of an associated headpost. MRI data allow optimizing positioning on the skull. The chamber is implanted into a precisely matched craniotomy, improving mechanical stability. For maximal biocompatibility, chamber and headpost are made out of titanium. It consists of an outer cylinder into which screws adjustably an inner one, called “well”, with a glass window on its bottom. This allows easy opening and closing whenever access to the cortex is needed, while preserving a sterile seal when closed. Moreover, simply rotating the well allows fine adjustment of the window’s distance with respect to the underlying cortex. Together with using a curved - rather than flat - glass window, this allows achieving gentle but continuous contact with the underlying cortex, which helps to delay tissue regrowth, notably of a neomembrane that must otherwise be surgically removed. The chamber’s sealing system combines a silicone elastomer and O-ring, minimizing infection risk and CSF leakage. Data from longitudinal two-photon imaging using genetically encoded fluorescent calcium sensors reveal high optical quality over months, with excellent resolution of neurons and their activity. We also demonstrate the feasibility of two-photon imaging of genetically encoded voltage sensors in the marmoset cortex in-vivo. Finally, although the chamber provides a durable, adaptable solution for long-term imaging studies in marmosets already in its current version, we discuss a few straightforward modifications that could likely improve its performance even further.

## 1. Introduction

Over the last decades, the marmoset (Callithrix Jacchus) has become an increasingly popular animal model in neuroscience research, especially when it comes to questions that concern specifically the primate brain (Mitchell and Leopold, 2015; Miller et al., 2016; D’Souza et al., 2021; Okano, 2021; Yamamori, 2021; Samandra et al., 2022). Optical imaging approaches, be they micro-(Sadakane et al., 2015a,b; Santisakultarm et al., 2016; Ebina et al., 2018; Pattadkal et al., 2024) or mesoscopic (Yamada et al., 2016; Song et al., 2022; Shimaoka et al., 2024; Song et al., 2024), have proven extremely valuable for acquiring such data in vivo, by recording from large neuronal populations at high spatiotemporal resolution, down to the millisecond and the individual cell level. Since these techniques need optical access to the location to be imaged, in our case the cortex, the marmoset’s lissencephaly comes particularly useful.

In order to gain optical access to the cortex, a craniotomy and a durectomy need to be performed, and a “chamber” needs to be implanted onto the skull in order to seal-off the brain from the external environment, which would inevitably result in infections and/or mechanical damage. Such chambers are of standard use in neuroscience applications allowing “chronic” recordings from the brain of the same animal over several months, and different versions have been developed in different laboratories, first for use in the macaque monkey (Shtoyerman et al., 2000; Chen et al., 2002) and, more recently, in smaller animals like the marmoset (Sadakane et al., 2015a,b; Santisakultarm et al., 2016; Yamada et al., 2016; Ebina et al., 2018; Paddadkal and Priebe, 2025s). Typically, they consist of a cylinder made out of a rigid biocompatible material (metal, nylon, glass…) and are fixed onto the skull in a way that provides both mechanical stability and a reliable seal, in most cases using bone-screws and/or dental- or bone-cement. Either the chamber itself is made out of glass, or it has a transparent glass (or plexiglass) window providing a seal towards the outside, through which the underlying cortex can be imaged. In designs where the window is positioned on the top of the chamber (Shtoyerman et al., 2000; Chen et al., 2002), the space between the glass and the brain (which, in some cases, is covered by an artificial transparent dura mater (Arieli et al., 2002)) is typically filled with agar, often containing some anti-inflammatory and/or antibiotic agents in order to prevent or topically treat eventual infections and/or inflammations (due to the large distance between glass and cortex, this latter design is problematic when used with typical microscopy objectives because of their short working distance). Finally, the transparent window is protected against mechanical shocks that may result in scratching or even breaking it by a metal lid screwed onto the chamber, which can be easily removed for imaging, and swiftly put back into place before returning the animal to its housing.

A major challenge of chronic optical imaging is posed by the regrowth of tissue on the cortical surface, as part of the healing process triggered by the trauma of durectomy. This tissue neo-membrane, which starts off very thin and fragile, thickens progressively and not only affects the optical quality of the images, but also impedes physical and chemical access to the cortical surface, as is needed, e.g., in order to stain the cortex with voltage sensitive dyes (Orbach et al., 1985; Shoham et al., 1999; Slovin et al., 2002). It begins to form with variable delays following dura resection, ranging from 3 weeks (Chen et al., 2002) to 4 months (Arieli et al., 2002) in the macaque, whereas in some of our marmosets it started to form as early as 2 weeks. Moreover, at least in our hands, removal of this neo-membrane in the marmoset was less successful than in the macaque, even when it had thickened enough to form a separate layer (Chen et al., 2002).

If the exact mechanisms underlying this tissue regrowth remain to be elucidated, it has been noticed that when the cortex is irritated or injured, this tissue regrowth is accelerated, whereas, when manipulating the cortex is limited to a minimum and it is kept in tight contact with an artificial dura mater (Arieli et al., 2002; Chen et al., 2002), regrowth is somewhat slowed down. In this spirit, Sadakane and colleagues (2015a) developed a “glass chamber”, where the exposed cortex is constantly in contact with a glass surface, the rational being that gentle but continuous pressure by a biocompatible, inert, smooth but rigid surface would slow down neo-membrane regrowth even more than the soft artificial dura mater used in previous studies. Concretely, Sadakane and colleagues’ (2015) glass chamber consisted of a few, circular, ~100µm thick microscope coverglasses, having the same diameter (3mm) as the (circular) craniotomy and assembled one on top of another with the help of an optical adhesive, such that the resulting transparent glass cylinder had a height slightly exceeding the depth of the craniotomy. Finally, a last, somewhat larger (5.5mm) coverglass was glued on top of this cylinder, forming a concentric rim of 1.25 mm around it. The resulting device was then inserted into the craniotomy somewhat like a “plug”, its lower glass surface touching the cortex and its upper rim touching the bone, to which it was permanently cemented in order to form a hermetic seal.

We tried this design ourselves in one animal; however, as it sometimes happens when durectomy takes longer than foreseen, the brain was slightly oedematic at the moment of sealing the implant onto the skull. When the cortex subsequently retracted, the contact with the plug’s lower glass surface was lost and so was the regrowth-inhibiting effect. This can in principle be avoided by piling up enough coverslips and “pushing back the cortex” to reach the level one estimates the cortex shall relax to once the oedema is reabsorbed. However, correctly estimating this level is difficult; moreover, excessive pressure can induce seizures or other health problems.

Another shortcoming of this design is that physical access to the cortex is lost once the implant is sealed, which not only makes it impossible to stain the cortex (required, e.g., for optical imaging of voltage sensitive fluorescent dyes), but it also prevents the use of topical antibiotics to treat eventual cortical infections. Last, we were somewhat worried with respect to the mechanical resistance of the implant because of the fragility of the top coverglass (~100µm thick).

In order to address these issues, we developed an original chamber design. It allows easy opening when needed, yet provides a sterile seal when closed. Moreover, it allows for easy adjustment of the position (depth) of the glass window with respect to the cortical surface, thus allowing it to accommodate an edematous brain when needed, and to preserve contact with the cortical surface when the brain retracts.

Recently, Pattadkal and co-workers (Pattadkal et al., 2025) published a chamber design somewhat similar to ours. Yet, despite their similarity, the two chambers differ under a number of aspects, most importantly: the size of the field of view, the range of depths to which the glass window can be adjusted, how the chambers anchor to the skull, and the usage of a curved rather than flat glass window. The advantages and disadvantages of the two designs are discussed in more detail in the “Discussion” section.

## 2. Materials and Methods

### 2.1 Tailoring the chamber and its position to each individual via anatomical MRI

Before chamber installation, each animal was anesthetized and an anatomical 3D MRI image of its head was obtained (Fig. 1) and used in order to choose the optimal position of the chamber and to fine-adjust the inclination of the “winglets” on the chamber’s external rim (see below), which would subsequently rest on the bone surrounding the craniotomy. In some particular positions, *e*.*g*., very occipital ones with the rim of the chamber above the occipital ridge (maximizing the amount of primary visual cortex in the field of view), some of the winglets were trimmed because they had no bone below them.

**FIGURE 1:**
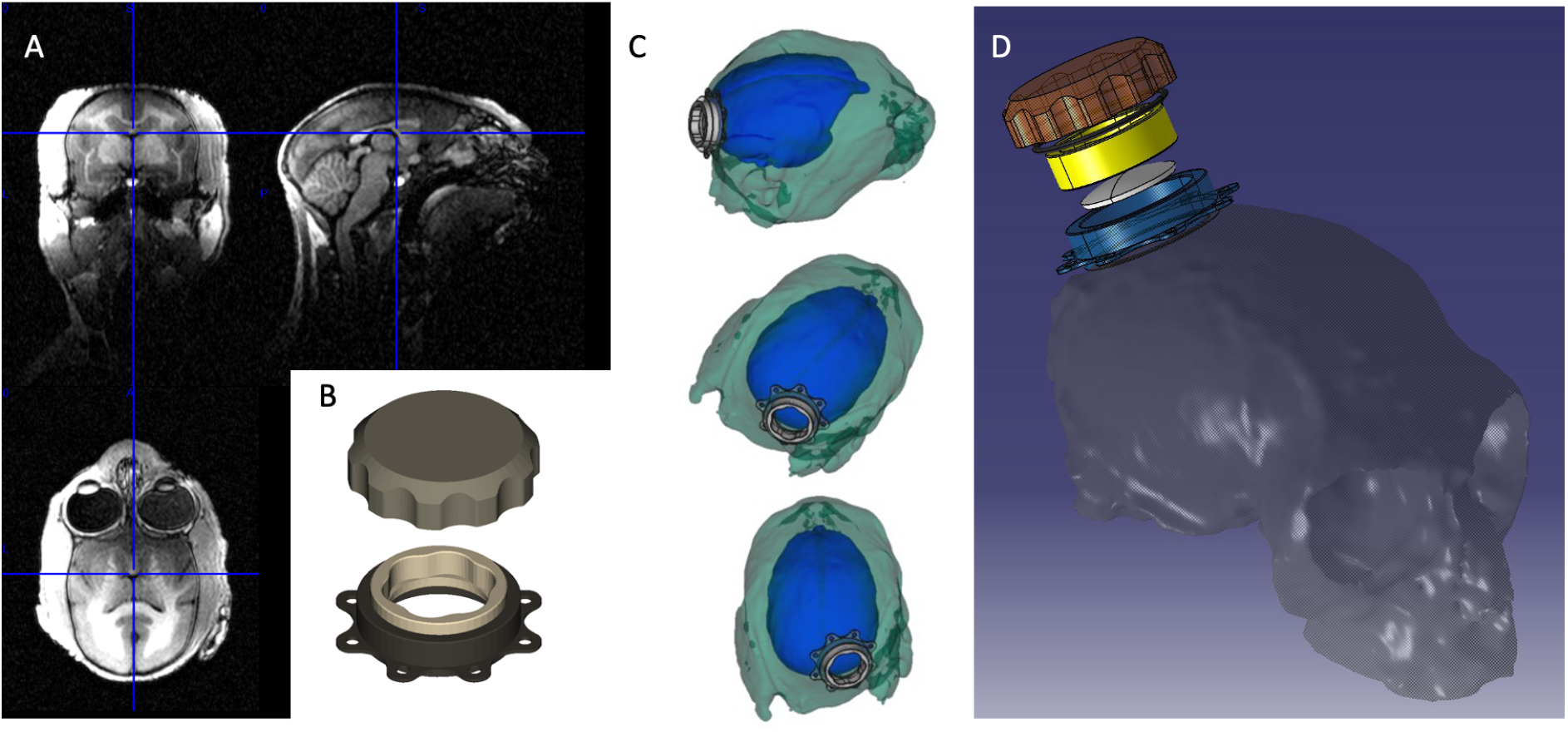
Fitting the chamber onto the skull; chamber parts. A: anatomical MRI scan (T2) allowing to optimally place the chamber and to bend its “winglets” to the skull’s curvature. B: Complete chamber and protective lid. C: 3D simulation of the chamber’s position on the skull, using the MRI segmentation of the bone and the brain (blue volume), allowing precise positioning of the chamber over the area of interest (here, V1). D: Disassembled view of chamber on skull. For the technical drawings see Supplementary Material

### 2.2 Chamber design

We first wondered whether it might not be possible to increase the amount of optically accessible area with respect to the 3 mm or 4 mm diameter design of, respectively, Sadakane et al. (2015a) and Pattadkal et al. (2025), because we wanted to perform mesoscopic wide field imaging in addition – and in complement – to two photon fluorescent microscopy. We found that a chamber with a clear aperture of 8 mm in diameter mounted onto a craniotomy of 11.5 mm is a good compromise between a large imageable area, component sizes that don’t raise excessive difficulties upon fabrication, and a physiologically well tolerated implant.

Our chamber (Fig. 2) consists of two main parts, namely two metal cylinders that screw one into the other. Starting from its lowest end for about 1mm - slightly less than the thickness of the bone - the outer cylinder has a smooth outer surface and an external diameter that exactly matches the craniotomy, into which it is to be inserted. A thin rim (visible in Fig. 2F) with 8 winglets follows, which can be adapted to the particular morphology of each individual by bending or cutting. This rim has a slightly larger diameter than the craniotomy, and functions as an end-stop together with the winglets, when the chamber is inserted into the craniotomy. This results in a tight seal when the chamber is finally cemented to the bone. The rest of this cylinder’s external wall is threaded up to its top, in order to allow a protective metal lid to be screwed onto it. The inner wall of this first, external, cylinder is threaded from top to bottom in order to allow the second, inner, cylinder, whose external wall is also threaded, to be screwed into it up to the desired depth. This inner cylinder has a notch on its bottom (visible in Fig. 2C,F), into which we glue a round coverglass using ultraviolet curing adhesive (NOA81; Norland Products Incorporated). Being thus closed at the bottom, we also call this inner cylinder “well” (or “piston”).

**FIGURE 2:**
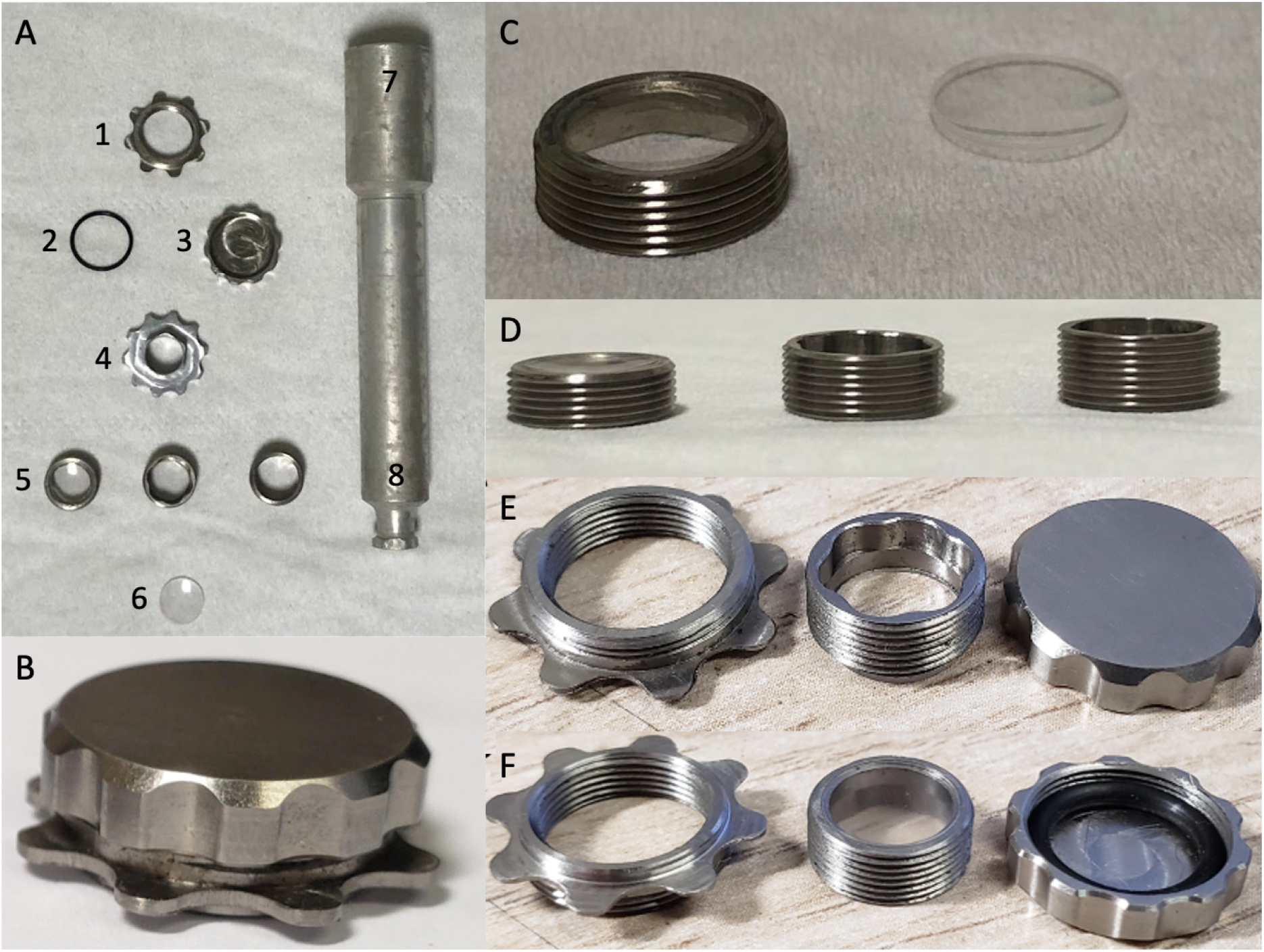
Chamber realisation. A: (1) external cylinder with small winglets that can be bent in order to optimally fit the skull; (2) rubber O-ring providing an airtight seal when the protective lid (3) is screwed onto the external cylinder (1). (4) Tool for screwing the inner cylinder (5) into the external one. Note the presence of a hole in its center, allowing to see the underlying vasculature through the coverglass and thus to assess the exact depth at which the two enter in physical contact. (5): Three inner cylinders (“wells”) of different length, see also (D), with mounted curved coverglasses. (6) Curved coverglass, which is normally glued onto the bottom of the well, provided with a receptive notch (see also C). (7,8) two-sided tool: its top end (7) fits onto the protective lid and allows quick and easy loosening and tightening of the latter. Its bottom end (8) allows easy positioning of the well within the external cylinder before fine-adjustment with key (4). B: Fully assembled chamber as it would be implanted upon the skull, with protective lid. C: Left, well with curved coverglass glued into the notch at its bottom (well is shown upside-down in the picture). Right, unmounted curved coverglass. D: side view of three wells of different length, allowing to adapt to the depth of the cortex within the craniotomy. E, from left to right: external cylinder, well and protective lid. The tools in A (4 and 8) fit into the 6 notches on the well’s inner wall. F same as E, but shown upside-down. Note the black rubber joint (O-ring) inside the protective lid.

Thanks to the described mechanism, the chamber can be easily opened and re-closed whenever needed (e.g., for cleaning, microinjections…). Most importantly, it is possible to (re-)adjust the inner cylinder’s depth within the outer cylinder simply by turning it: this allows to make sure the coverglass on its bottom stays in contact with the cortex even following eventual changes in the brain’s volume. In order to ensure biocompatibility, all metal parts are made out of titanium.

We found it useful to screw the well into the external cylinder under visual guidance. Indeed, doing so while observing the blood vessel pattern at the bottom of the chamber allows to position the coverglass such that it does indeed touch the cortical surface, yet without exerting excessive pressure: one simply has to stop turning as soon as the vessels are seen to be dragged by the coverglass, somewhat following the well’s rotation. The reason for the central hole in the center of the small tool (#4) in Fig. 2A is precisely to allow this observation.

The chamber is complemented by a headpost (Fig. 3) allowing to swiftly and reproducibly fixate the animal’s head. While headfixation via a headpost is a necessary condition while imaging in the awake animal due to the much better comfort they provide to the animal as opposed to earbars, we also found the presence of a headpost useful in the anesthetized condition, because of its much easier mode of operation as compared to earbars, the correct positioning of which can be tricky.

**FIGURE 3:**
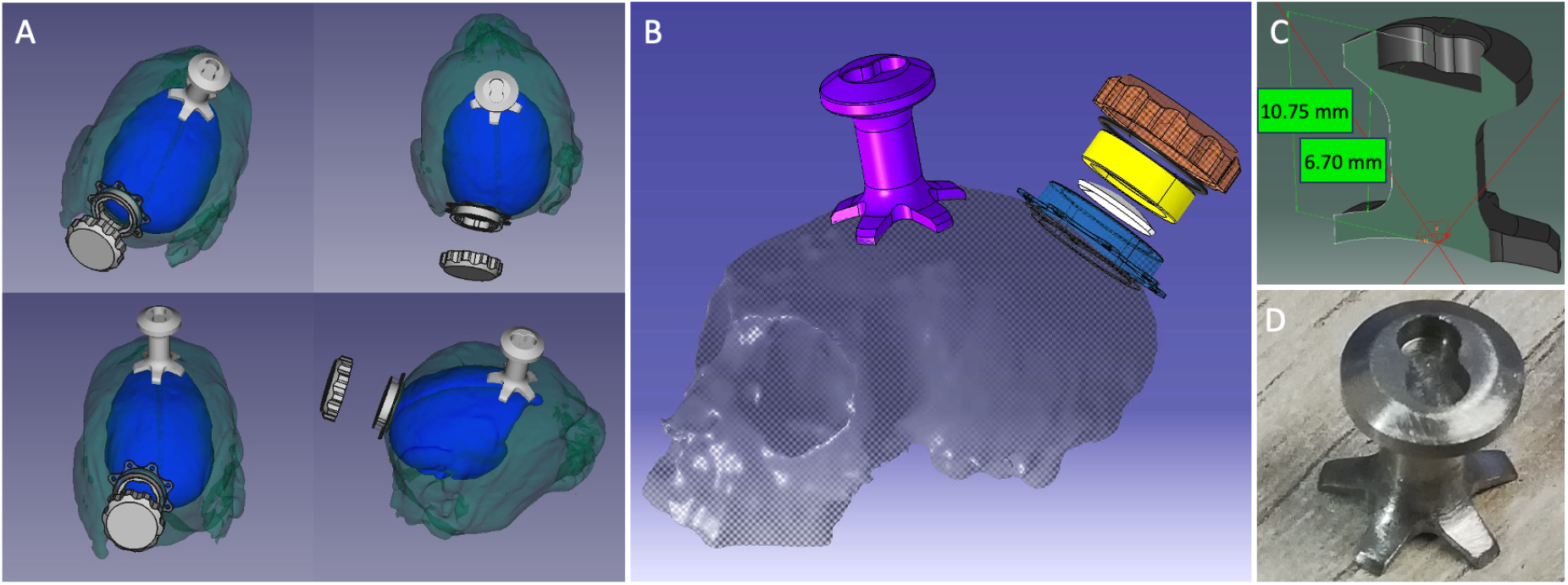
Headpost. A: Four views of the chamber with additional headpost, projected onto a marmoset skull segmented from an anatomical MRI scan (color codes as in Fig 1C). B: detailed view from the side. C,D: Section with principal measurements and photo of headpost. For details see the technical drawings in the Supplementary Material

### 2.3 Large changes of brain volume, optimization of contact surface

For the sake of compatibility with the small working distance of most microscope objectives, the whole chamber was kept as flat as possible (total height of external cylinder, without protective lid: 3.30 mm). As a result, the range of depths within which a well of a given length can be adjusted is quite limited and, in some cases, not enough: *e*.*g*., in case of strong oedema, a long well will protrude beyond the outer cylinder’s external edge impeding the protective lid to be screwed onto it. Conversely, in the case of a hypovolumic brain, a short well will not be able to go deep enough to preserve contact with the cortex. In order to deal with this problem, wells of different length were machined, and each time the chamber was opened, at least three sterilized ones were kept ready for use. Fig. 2D shows an example of wells being 3, 4, and 5 mm long (other sizes are obviously possible).

Finally, in order to optimize the contact between the glass surface and the cortex, we used a curved rather than a flat glass on the bottom of the well (8 mm diameter, 500 µm thickness), more precisely, a spherical lid having the radius of curvature of 11 mm that we had obtained as average curvature of the marmoset brain on the area of interest (V1) measured from the anatomical MRI data obtained from 6 marmosets.

### 2.4 Sealing

A perfect sealing of the chamber is of extreme importance in order to protect the cortex from infections on the one hand, and, on the other, to avoid the cerebrospinal fluid (CSF) to leak out of the chamber thus resulting in a loss of intracranial pressure. A first barrier against the external environment is provided by the thread between the well and the internal wall of the external cylinder. However, rather than aiming for a tight fit between the two cylinders that would be challenging to machine, we decided to leave some clearance between them and to provide the seal by injecting a silicone elastomer (“Kwik-Sil”; World Precision Instruments) into the gap between the two parts. It is applied as a liquid – thus efficiently penetrating into the thread by capillarity – and solidifies within a few minutes giving rise to an elastic joint that not only perfectly adheres to the metal, but which can also easily be removed and renewed at every chamber opening.

A second barrier is provided by a rubber O-ring fixed to the inside of the protective titanium lid that is screwed onto the outer cylinder while the animal is in the animal facility, and which is removed during imaging. This joint provides a seal between the lid’s ceiling and the top of the chamber’s external cylinder. As already mentioned, for this seal to be effective the well must not extend beyond the edge of the chamber, otherwise it can prevent the lid to reach its optimal position, possibly disrupting the contact of the joint with the chamber rim. Even worse, if the well is in contact with the lid or its joint, turning the lid may also drag the well with it, compromising the sealing provided by the silicon elastomer and opening the door for infections. This risk was one of the main reasons for always having wells of different length available.

### 2.5 Animals and ethics

Four animals, 1 male and 3 females aged between 17 and 70 months (young adults), were used to test the new imaging chamber. The animals were carefully monitored, especially during the few hours after the implantation surgery and for several days after each procedure to ensure their well-being. A specialized welfare assessment grid for marmosets was developed in our laboratory to evaluate their health condition following surgeries. This grid allows scoring of various parameters to identify specific symptoms, enabling tailored post-operative treatment and care for each animal based on its individual needs. The protocol was pre-approved by the local ethics committee (CEEA71) and authorized by the Ministry of Research and Higher Education according to EU Directive 2010/63 for the protection of animals used for scientific purposes.

### 2.6 Anaesthesia and physiological parameters

Induction: For both recording sessions and surgical procedures, the animal’s anesthesia was consistently induced following the same protocol. Animals were deprived of solid food at least four hours prior to anesthesia induction and water was removed from the cage at least 45 minutes before. Following the capture of the animal, an initial intramuscular injection of glycopyrrolate (0.01 mg/kg) was administered into the thigh to minimize salivary secretions, prevent bradycardia and any risk of choking during anesthesia induction. Twenty minutes later, a second intramuscular injection of alfaxalone (12 mg/kg) and midazolam (0.1 mg/kg) was administered in order to induce anesthesia. This anesthetic/sedative combination proved effective in marmosets, reliably inducing narcosis and rapid muscle tone loss, typically observed within 5 to 10 minutes after injection.

Chamber implantation surgery: after induction, anesthesia was initially maintained via inhalation of sevoflurane (0.5–3%) in 100% oxygen, delivered through a marmoset-adapted breathing mask, for the time needed to set up the animal. An intravenous 26G catheter was then placed in the saphenous vein (located at the back of the calf), with the purpose of administering intravenously NaCl 0,9% (5 ml/kg/h) to keep the animal hydrated and to prevent hypotension, and, at constant rate, alfaxalone (10–15 mg/kg/h) to maintain anesthesia, narcosis and muscle relaxation, and fentanyl (10 µg/kg/h) to provide analgesia. Sevoflurane was stopped after infusion started, although later, during surgery and depending on the anesthesia level, sevoflurane at low concentrations could be reintroduced in addition to the infusion. After a bolus of alfaxalone (3 mg/kg), the animal was intubated with a custom made probe and respiration was controlled using a specialized small-animal respirator. It delivered a gas mixture of 70% O_2_ and 30% air at a rate of 1 L/min and a respiratory rate was fitted to target end tidal CO2 levels between 35-45 mmHg in volume-controlled mode. The fentanyl infusion was stopped 30 min before the end of the anesthesia to prevent any further respiratory problem.

Viral injections were performed after a two-weeks recovery period following the chamber implantation surgery. After induction of anesthesia as described above, it was maintained using isoflurane (1–3.5%) or sevoflurane (0.5–3%) in 100% O_2_, administered via inhalation through a custom made 3D stereotaxic mask. The same anesthesia protocol was used for recording sessions, except that the isoflurane level was gradually lowered during the session in order to reach the least amount needed to keep the animal anesthetized.

During all procedures, the animal’s vital signs were closely monitored. This included heart rate, oxygen saturation (SpO_2_) and body temperature. Blood pressure and end-tidal CO2 when animals were intubated are also monitored during surgical procedures. The target physiological parameters were as follows: heart rate of 170–230 bpm, SpO_2_ at 100%, body temperature maintained between 37.5°C and 38.5°C, and blood pressure around 120/80 mmHg. The anesthesia level was dynamically adjusted as needed to ensure the animal’s stability throughout the procedure.

### 2.7 Chamber implantation surgery

After anesthesia induction, the animals were prepared for the procedure. The hair was shaved from the head, the back of the thighs and the limbs, to accommodate various monitoring sensors. Subcutaneous injections of ropivacaine were administered at the planned incision site (0,2 mg per site) and at three different points inside the ear canal (0,1 mg for each site) to prevent discomfort from the ear bars during surgery according to good practice in anesthesia for eye-nose-throat surgeries, and a subcutaneous injection of meloxicam (0,2 mg/kg) was administered to prevent any other pain and inflammation. After the intravenous lines were placed and the animal was intubated, it was positioned in a stereotaxic frame. The scalp was then disinfected with alternating applications of povidone-iodine and ethanol. Sterile drapes (we used 3M™ Loban™ Antimicrobial Incise drapes that adhere securely to the skin thus reducing the risk of surgical site infections by immobilising bacteria and providing continuous antimicrobial infusion) were applied to cover the animal and non sterile instruments, and the surgeon wore sterile protective clothing and gloves throughout the procedure.

In the following, we describe the surgical procedure for a chamber installation on the left hemisphere. A coronal skin incision was performed in front of the ears. We preserved the skin, the subcutaneous tissue and the periost by applying hydrated compresses in order to facilitate the healing process. We partially removed the occipital part of the left temporalis muscle with an elevator to preserve the fascia for the healing process. The attachments of the right muscle were retracted only if required in order to be able to perform the next steps. When access to the occipital ridge was necessary in order to position the chamber as posterior as possible for optimal V1 visibility, we also needed to slightly retract the obliquus and rectus capitis muscles.

The craniotomy was then performed using a homemade trephine bone saw developed in our laboratory’s mechanical workshop. This tool consists of a stainless steel cylinder with sharp teeth mounted on a drill compatible with our micromotor (Figure 4A) and allows for a craniotomy precisely matching the outer diameter of the lower part of our imaging chamber, that is, 11.5 mm. Once the circular bone lid cut by the trephine was removed, the contours were smoothened by removing remaining thin parts of the bone to avoid injuring the underlying tissues upon contact. If (and only if) necessary, the craniotomy was slightly enlarged using a standard drill, the goal being to be able to place the chamber into the craniotomy with a fit as perfect as possible, which was achieved by repeated testing. Indeed, any gaps at its border would constitute potential entry points for infections.

**FIGURE 4:**
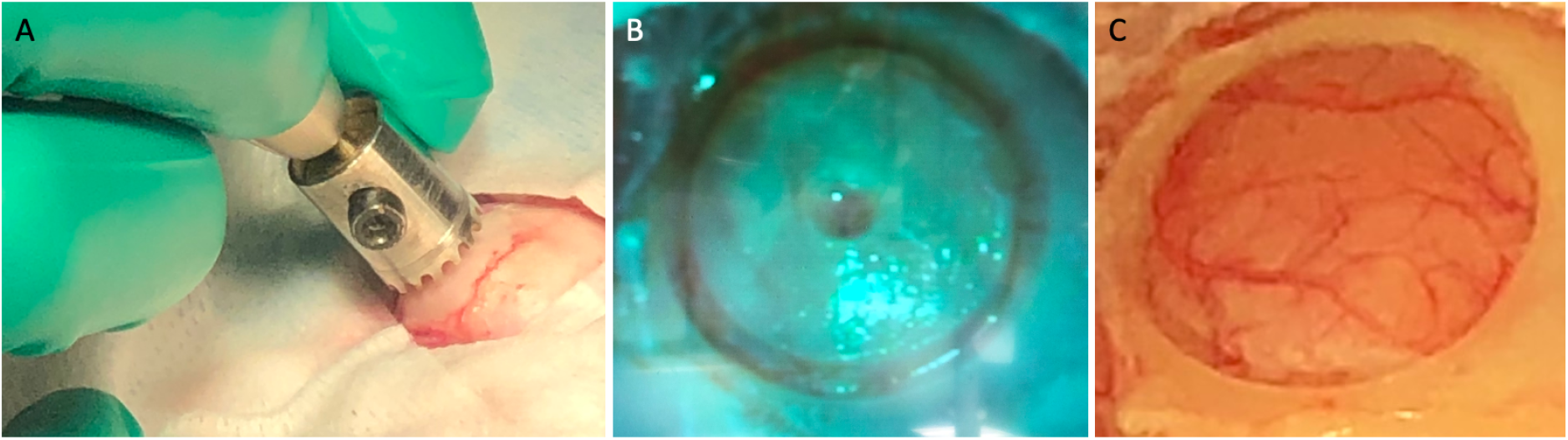
Craniotomy. A: A homemade trephine bone saw mounted on an electric drill with controllable speed, torque, light and fluid irrigation (micromotor for surgery NSK, surgic XT plus). Moist gauze pads cover the bone in order to prevent it from drying; only the surface that is strictly necessary for the operation is exposed to the air. B: Nearly complete craniotomy. Note dural vessels visible through the much thinned bone at a few locations in the groove. In order to avoid blessing the dura mater, the very last layer of bone is cut using a drill with a 0.5mm diamond burr. C: Completed craniotomy, before dura opening. Note the regularity of the border, allowing a clean and tight insertion of the chamber.

Once we were satisfied with the craniotomy, the chamber was put aside in order to leave space for the durectomy. Importantly, in order to prevent brain oedema following the dural opening, mannitol (1 g/kg, complemented by additional 1 g/kg in case of persisting oedema) had been administered intravenously for some 20 minutes while refining the craniotomy, and the respiration rate was increased to induce hyperventilation targeted to etCO2 28-32 mmHg.

We found it practical to begin the durectomy by making an initial incision in the dura mater using a 30G needle. The dural opening was then continued using a “dura forceps”, that is, tweezers with a very fine tip, provided with rough inner surfaces that facilitate to hold the dura mater (Dumont ultra fine tweezers) and micro-scissors, approaching the edge of the craniotomy as closely as possible (Fig. 5). Once the dural opening was completed, 2-4 screws (Universal self-tapping screws, 1.2 mm diameter x 2 mm or 3 mm length (OSTEOMED, France) were inserted into the bone around the craniotomy. These screws, embedded in the dental resin used to secure the chamber to the skull, provided additional stability to the fixation, which the resin alone could not achieve. A primer (Peak Universal Bond, Ultradent) was then applied on the skull, all around the craniotomy and screws, and was first photopolymerized using an ultraviolet lamp. Thereafter, dental resin (LC Block-Out, Ultradent) was applied over the chamber’s winglets and around the chamber, covering the screws. The resin was then photopolymerized a second time to securely fix the entire assembly to the skull. Once everything was secured, the well was screwed into the inner thread of the chamber and lowered to its maximum extent, ensuring that the glass attached to its tip made contact with the cortex without applying any pressure on the brain.

**FIGURE 5:**
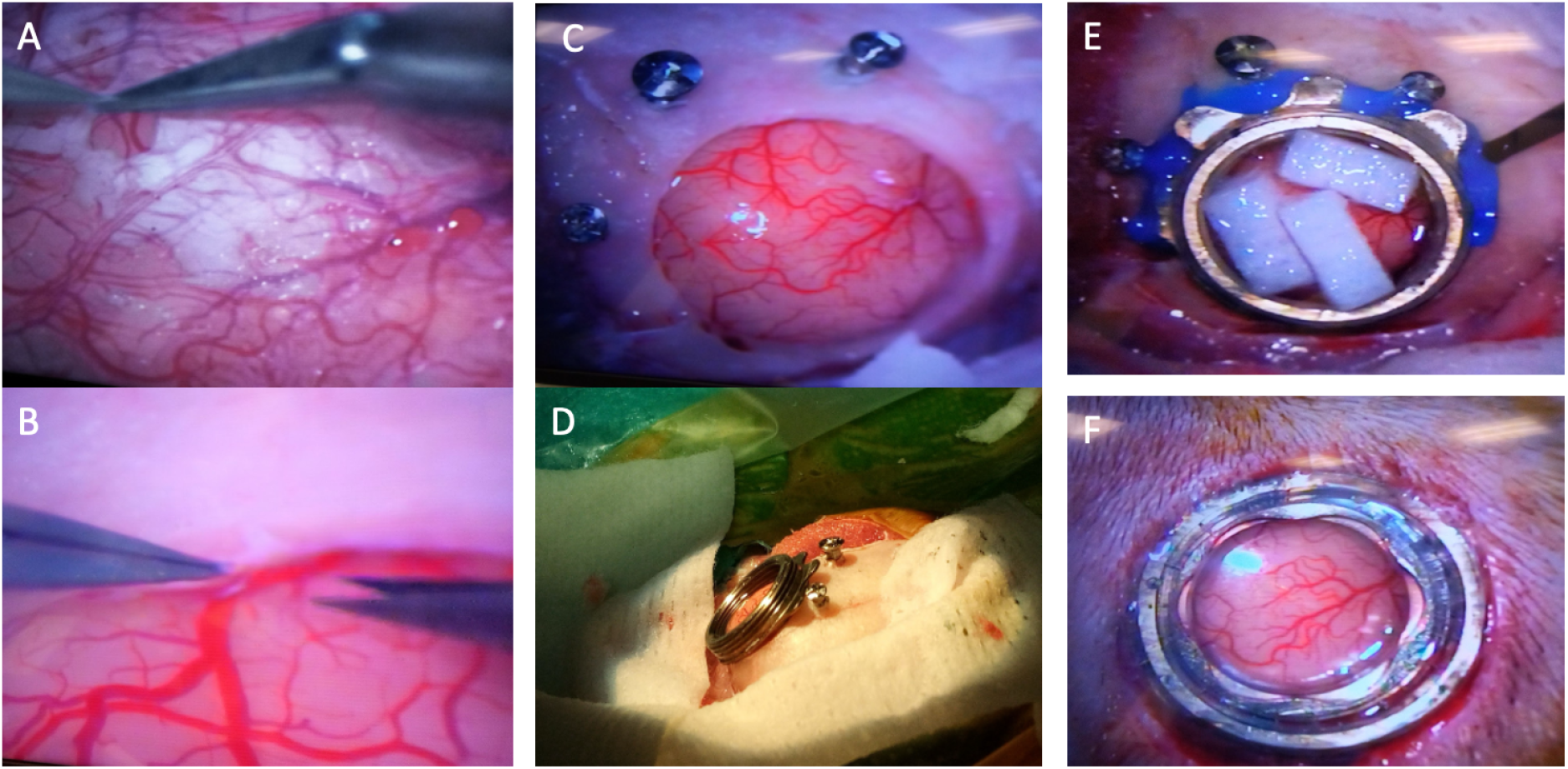
Excision of dura mater (durectomy) and chamber fixation. A: After a “pinch” using a 30G needle, the dura mater is carefully removed using miniature scissors, approaching as closely as possible the edge of the craniotomy (B). C, D: Screws placed around the craniotomy (before pinching the dura) provide anchoring points in order to increase the chamber’s mechanical stability. Note that the chamber being positioned very close to the occipital ridge, some winglets had no bone to adhere to and have thus been clipped off (at the left in the image, visible also at the bottom in (E)). E: The screws and the imaging chamber are covered with photopolymerizable dental resin (blue). During photo-polimerization, which requires shining ultraviolet light onto the resin for a few minutes, the brain is protected by small pieces of wet neuro-pads. F: The chamber is closed with a well with a curved coverglass on its bottom that touches the cortex. Note the tight adherence of the skin to the chamber (see also Fig. 6B)

After securing the chamber, we moved on to implanting the headpost. Five screws were inserted into the frontal region of the skull to allow for the attachment of the head post (Fig. 6A). The headpost, which has 5 feet on its bottom (Fig. 3C,D), was positioned at the center of these screws and secured using dental cement (Palacos R+G, Heraus Medical). The skin was then repositioned and sutured such that it completely covered the two implants. Finally, two new, tight, incisions were made in order to allow the imaging chamber and the headpost to emerge from under the skin (Fig. 6B). As compared to suturing around the implants, this technique promotes better healing of the initial incision and overall cicatrization, while minimizing the risks of complications related to inflammation and scarring. In our case, we implanted the headpost and the chamber in one single surgery, but they can also be implanted in two separate surgeries. This latter choice can be preferable if the animal needs to undergo many head-fixed training sessions before starting with imaging, or simply to get the animal to accept to be head-fixed for imaging if the latter happens in the awake state: obviously, one strives to perform craniotomy and durectomy as late as possible in order to minimize the risk of infections, tissue regrowth etc. (see Figs. 10 and 11).

**FIGURE 6:**
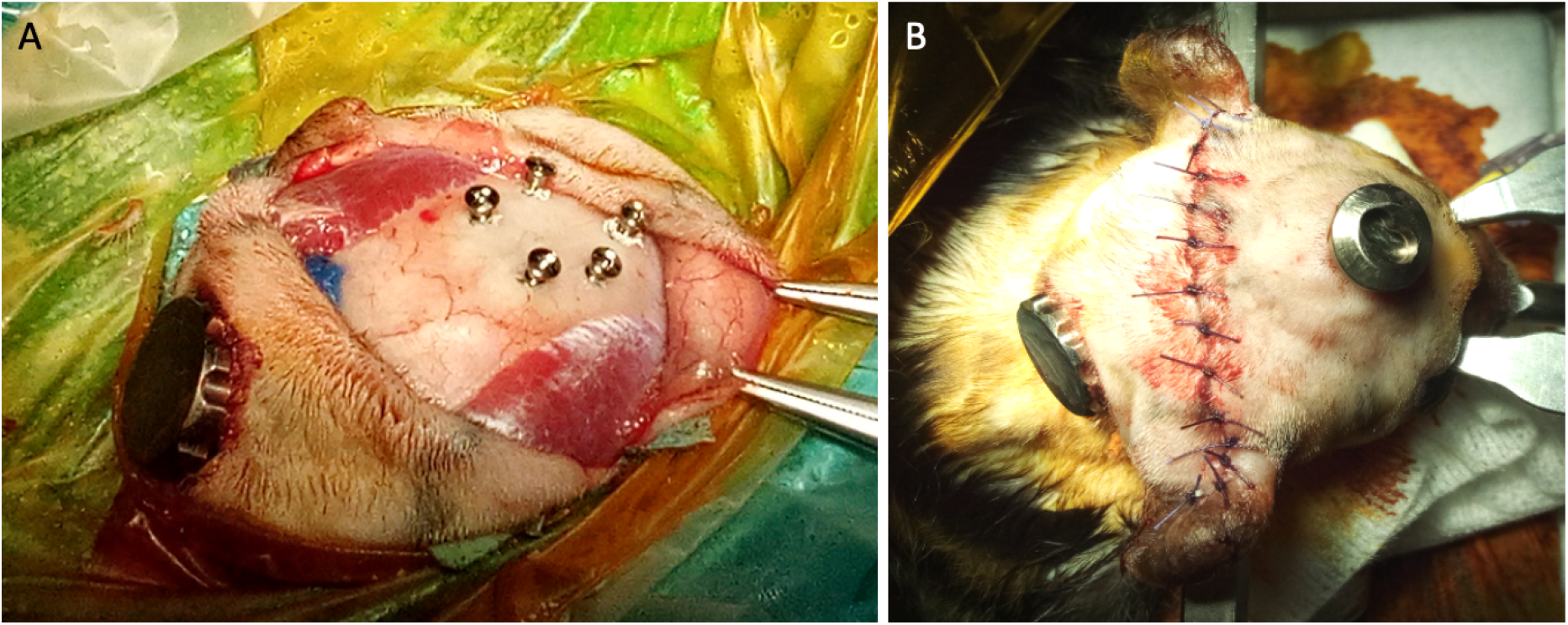
Headpost. A: Placement of the screws to fix the headpost. B: Imaging chamber, headpost and sutures at the end of the implantation surgery. Note the tight adherence of the skin to the implants.

Anesthesia was then stopped to awaken the animal. Meloxicam (0.2 mg/kg/day) was administered for three to six days, and buprenorphine (10 µg/kg/day) was given for at least one to three days to manage both inflammation and pain. Antibiotic coverage was provided through the administration of long-acting oxytetracycline (20 mg/kg/72h) during three to six days. The duration of the various treatments could vary based on each animal’s well-being score following the surgery.

### 2.8 Microinjections

A minimum of two weeks after the implantation surgery, the animal was anesthetized again in order to perform the transfection with our GeCI-AAV constructs, that is, AAV9-mThy1-tTA at a titre of 5.66.10^12^ particles/µL mixed in equal parts with AAV9-Tre3-GCaMP7f at a titre of 7.93.10^12^ particles/µL, or AAV9-hSyn-Ribo-jGCaMP8m at a titre of 5.5.10^12^ particles/µL. One animal was transfected with a GeVI,, Jedi-2P, that is, AAV2/9-hSyn-JEDI-2P-WPRE at a titre of 2 × 10^13^ particles/µL.

These compounds were injected into the cortex using a glass micropipette and a precision microinjector (Nanoject III, Drummond). We chose to use this microinjector because it allows to reliably inject precise volumes at very slow injection rates, which, together with using micropipettes with very fine tips, helps minimizing cortical damage during viral injections, which is crucial for the quality of the subsequent imaging sessions. The micropipettes we used consisted of pulled glass capillaries with tips having an external diameter of 30 to 60 µm; they were beveled at an angle of 45° in order to penetrate cortical tissue without breaking, and sterilised before usage (Fig. 7A). For injection volume, speed etc., see “Technical Details”.

**FIGURE 7:**
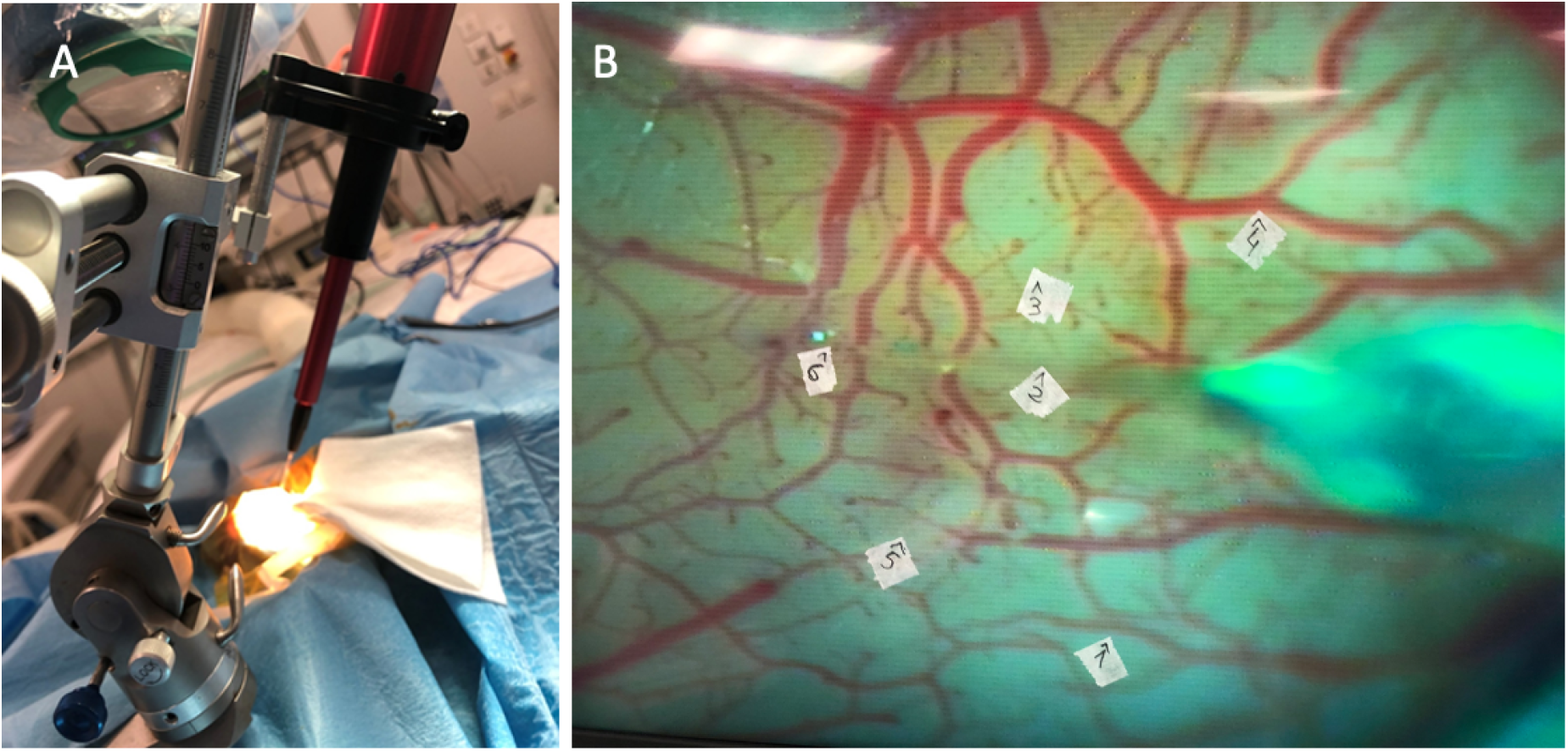
Microinjections for transfection. A: Placement of the motorized microinjector in the stereotaxic frame above the head of the animal (overexposed in the image due to reflections from the edge of the chamber), which is installed in the same stereotaxic frame. B: Six locations for injections on the cortex have been selected under the surgical microscope and marked with small pieces of auto-adhesive paper on the computer screen that displays the image provided by the camera connected to the surgical microscope’s imaging port. Note the tip of the glass micropipette (its body is visible out of focus on the right) inserted into the cortex at location #6.

After anesthesia induction and preparation identical to that of the implantation surgery, the chamber was reopened in a sterile environment. Before reopening the chamber, mannitol was administered intravenously (2g/kg during 20 minutes) to minimize oedema through the open chamber. The chamber was then opened by removing the well and three to six injection sites were visually selected with the help of a surgical microscope between the different blood vessels on the visual cortex, as posterior as possible, in order to target V1, the primary visual cortex (Fig 7B). The injector was then prepared and placed on the stereotaxic frame. Different injection parameters were tested to determine the optimal conditions for transfecting as many cells as possible yet avoiding overexpression, mechanical damage and leaking out of the injection site, keeping in mind that it is also important to minimize the time the cortical surface remains exposed in order to avoid oedema.

Matching these requirements implies finding the optimal combination of volume par injection site, number of sites, injection rate, titre, and depth. We tested two depths below the cortical surface (500 µm and 700 µm), three injection rates (60 nl/min, 240 nl/min, and 600 nl/min), two injection volumes (500 nl or 1 µL per injection site) and up to 8 injections but keeping the total volume injected below 5 µL per animal. Even at the highest injection rate the transfection worked satisfactorily, also because we created some space in the tissue for the liquid to be injected by advancing the micropipette some 50 µm beyond its desired depth and retracting it back the same distance before starting the injection. Once the injection finished, the pipette was left in place for 5 minutes before extracting it, thus providing a sort of plug and getting the injected liquid to diffuse into the tissue rather than to leak out along the pipette’s insertion channel. Once all the injections were completed, the chamber was closed with the well, and the thread was resealed using Kwik-Sil (World Precision Instruments) before screwing on the protective lid and waking the animal. Meloxicam (0.2 mg/kg/day) and oxytetracycline (20 mg/kg/72 hours) were administered to prevent pain, inflammation, or infection caused by the injections.

## 3. Results

### 3.1 Recording quality – 2p microscopy: GeCIs

We transfected several animals with a genetically encoded intracellular calcium concentration indicator (GeCI), JCaMP7 linked to a TET (tetracycline)-inducible system for AAV. Two components of the TET-Off system were cloned into separate AAV vectors: (i) the tetracycline-controlled transactivator (tTA) under the control of the Thy1S promoter (a modified version of Thy1 promoter), and (ii) GCaMP7f under the control of the tetracycline response element (TRE3) promoter. This allows to control the expression of the calcium indicator by administration of the antibiotic Doxycycline, e.g., in the drinking water of the animal (Sadakane., 2015). In the absence of Doxycycline (Dox), tTA constitutively activates expression of a transgene under the TRE3 promoter, whereas presence of Dox prevents the binding of the tTA to the TRE3 and inhibits transgene expression. The details on how the microinjections were performed have been described in the previous section.

The data shown in Figure 9 show GCaMP7f injected regions of V1, imaged in the same animal, at different timepoints after transfection.

The cortex’ imageability remained excellent over more than 5 months after dura opening, including a break of about one month during which Doxycycline had been administered to the animal (in the drinking water) in order to depress the expression of GCaMP7. Consistently with the results reported by Sadakane and colleagues (2015), after Doxycycline administration was stopped the expression returned to previous levels (Fig. 8, E,F).

**FIGURE 8:**
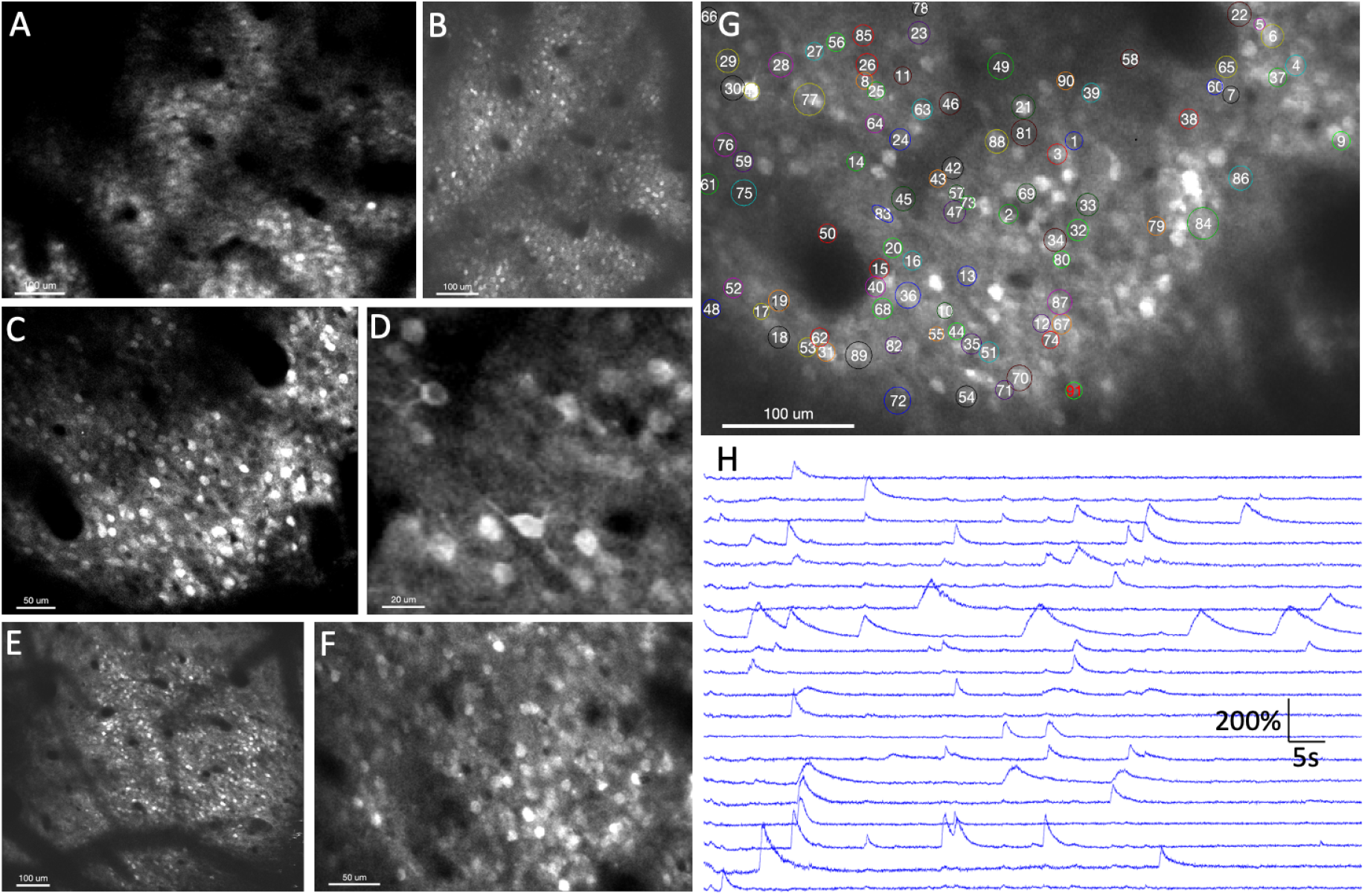
Chronic imaging of GeCIs. A-F: Crisp fluorescence images of GCaMP7-transfected neurons were obtained with a two photon microscope in galvo-resonant mode (31Hz) in the primary visual cortex of an anesthetized marmoset. The images were taken at 5 (A), 8 (B), 9 (C,D) and 22 weeks (E,F) after craniotomy and dura opening (which was followed two weeks later by transfection via microinjections). (E,F) were obtained about 4 weeks after having stopped Doxycycline treatment D) and (F) are a zoom into (C) and (E), respectively: note the high optical quality, allowing to clearly resolve some dendrites and cellular nuclei in (D). Images are time-averages (106 - 160ms). G-H: during the same session as in (C), we also recorded a 1.5 minutes long movie of the cells’ spontaneous activity (framerate: 47 Hz). 91 of the most clearly active cells were selected by hand (G) and their fluorescence time courses calculated. Panel (H) shows 20 representative examples of those (chosen out of the 91 for clarity of display), after subtraction of background and normalization by each time course’s temporal mean (after subtraction of 0.7 times the mean of the background). The upwards deflections of the traces correspond to an increase in fluorescence due to intracellular Calcium increase resulting from one or (in most cases) several fired action potentials.

As expected, neuronal activity - spontaneous in our case - depended strongly on the depth of anesthesia. When the latter was light, we could observe typical patterns of activity, in hundreds of cells. After re-registration of all frames in order to correct for small movements, cells were hand-selected and their raw fluorescence time course was bleaching-corrected and background-subtracted. Finally, each resulting trace was normalized by its temporal mean (90s). Figure 8H shows 20 representative examples of such time courses.

### 3.2 Recording quality – 2p microscopy: GEVIs

We also transfected animals with a genetically encoded membrane potential indicator, Jedi-2p (Liou et al., 2022). The data shown in Figure 9 were obtained in an anesthetized marmoset 1 month after transfection. The high-resolution image in Fig. 9A was taken in galvanometric scanning mode with relatively long dwell time (~10ms). It shows that the fluorescence is concentrated in the cell-membranes, confirming a correct localization of the indicator in the sample. For functional imaging we switched to resonant scanning, using our microscope’s second scanning head. Due to the much shorter dwell time the signal-to noise was considerably reduced (Fig. 9B), yet individual cells could still be distinguished.

We thus moved the objective to a position where a few clearly visible cells were clustering in the middle of the image (Fig. 9B, arrows), where SNR is optimal. In order to record the fast changes in fluorescence of the Jedi-2P indicator resulting from changes in membrane potential (action potentials), one obviously needs a much higher framerate than that obtainable by full-frame imaging (31Hz). This was achieved by reducing the number of scanned lines to a narrow stripe of only 16 (out of 512) lines, centered on the previously identified cells, resulting in a scan rate of 990 Hz. This stripe, averaged over 90s, is shown in overlay on the whole-frame in Fig. 9B.

**FIGURE 9:**
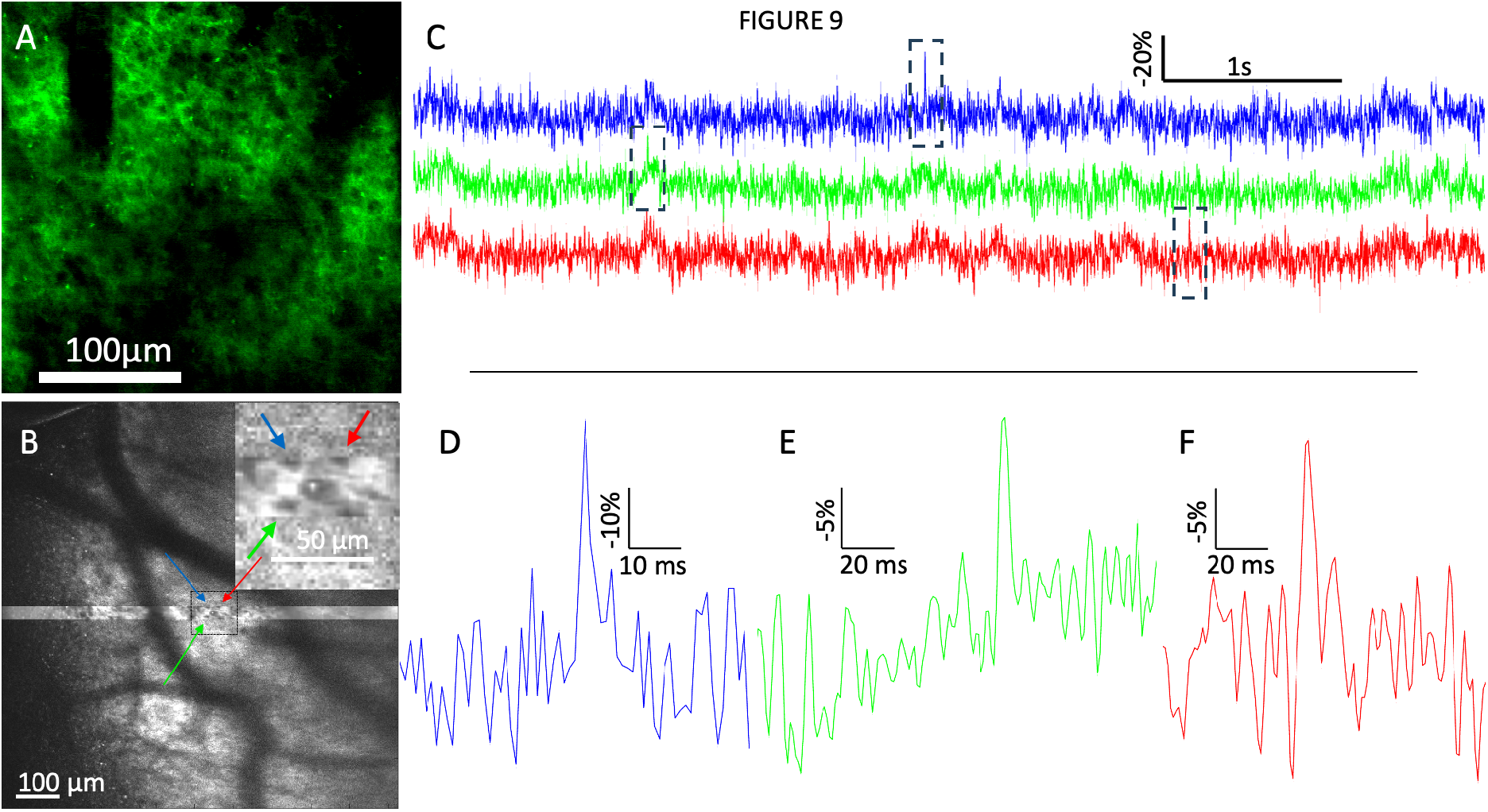
Imaging Jedi-2p-transfected neurons in the anesthetized marmoset primary visual cortex, in the absence of visual stimulation, one month after transfection. A: 2-photon fluorescence image obtained in slow raster scanning mode. B: 2-photon fluorescence image obtained at 31 Hz in full-field resonant scanning mode. Note that the imaged region in (B) is not the same as in (A). A thin stripe (16 pixels) was selected centered on some of the most clearly visible cells in the center of the image, averaged over time (35s) in order to increase the SNR, and shown as an overlay in B. The arrows indicate 3 of the most clearly visible cells. Inset: zoom into the region within the dashed square. C: time courses of the bleaching-corrected and background-subtracted fluorescence, normalized by its temporal mean (35s), recorded from the 3 cells shown by arrows in (B) (same color code). D-F: zoom into the regions outlined by the dashed rectangles in (C). Time courses in (C), (E) and (F) (but not in (D)) were temporally smoothed by convolution with a Gaussian of 5 ms standard deviation.

Although lacking confirmation by the gold-standard of simultaneous microelectrode recording, and despite a much lower SNR than that obtainable when imaging with dynamic holograms (ULoVE) (Villette et al., 2019, Liou et al., 2022), the events shown in (Fig. 9C-F), are clearly compatible with changes in fluorescence resulting from individual action potentials, both in amplitude and in duration (Liou et al., 2022). When moving to less bright cells or away from the center of the field of view, spike detection became obviously much more difficult, as expected from the increase in the level of (photonic) noise.

### 3.4 Imaging quality deterioration: tissue regrowth

As it has has been extensively documented previously (Arieli et al., 2002; Paddatkal et al., 2025), also in our case tissue regrowth set in at variable delays following durectomy, notably in the form of an initially very thin and fragile neomembrane, which, however, thickens with time and is progressively colonized by neovessels (Fig. 10).

**FIGURE 10:**
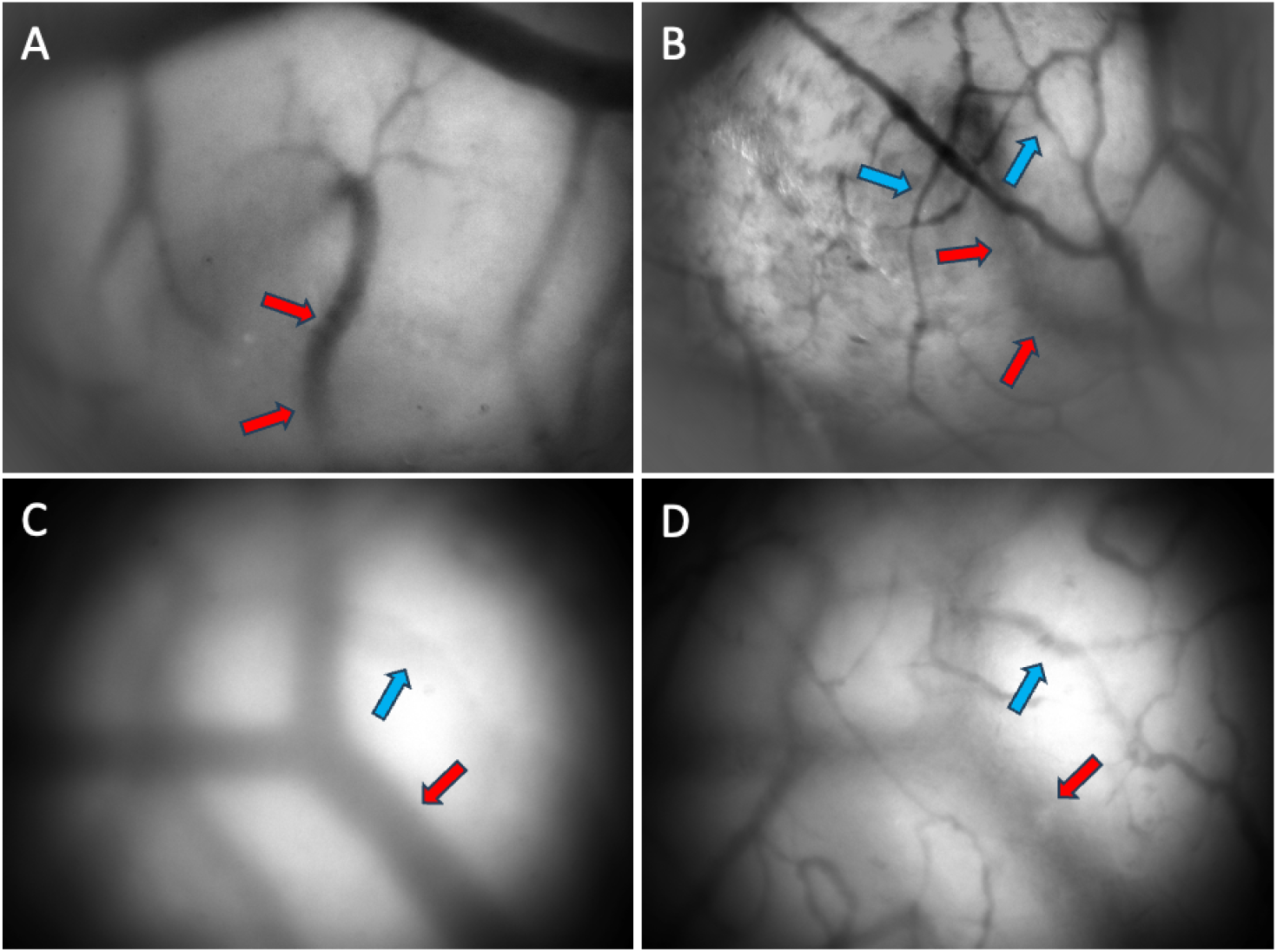
Example of a marmoset where neomembrane regrowth started at about 5 weeks after dura removal. A: image of cortex at 5 weeks after dura removal, showing a relatively clear image of the cortical vasculature despite a slight blur due to the onset of neomembrane regrowth. B: same as (A), but one week later (camera was slightly rotated between the two sessions), showing a layer of neovessels (blue arrows) that had grown above the cortical vasculature (red arrows, pointing to the same vessel in the two images) can still somewhat be distinguished through the neomembrane. C: Another cortical location on the same animal, 8 weeks after dura removal: despite focus on cortical vasculature (red arrow), its image is blurred due to light scattering by the neomembrane. An out-of-focus neo-vessel is also visible (blue arrow) in the image. D: same as (C), but focussing on the neomembrane, i.e., some 150 µm above the focus in (C). As a consequence, the neovessels (blue arrow) appear as sharp, whereas the cortical vasculature (red arrow) doesn’t.

This neomembrane regrowth was not gradual and its onset was different from one individual to another: in some cases it began as soon as one month after durectomy, whereas in others the cortex stayed clear for several months with good to excellent optical quality, yielding clear and crisp images upon two-photon microscopy up to 5 months after dura opening (Fig. 8). However, once the regrowth started (visible by a light blur in Fig. 10A), it advanced very quickly in all cases, thus rapidly deteriorating the optical quality of the preparation (note the vascularization of the neomembrane clearly visible in Fig. 10B, only one week after the image shown in Fig. 10A). In that particular animal, cellular resolution was completely lost as early as two months after dura removal, due to strong neomembrane formation. We hypothesize that this rapid regrowth was likely due to a lack of adherence of the coverglass - which in that particular animal was still flat - to the cortex. To maximize this adherence, we thus switched from flat to curved coverglasses matching the cortex’ curvature, which allowed slowing down regrowth by at least a factor of 2.

When the neomembrane had reached a thickness hampering cellular resolution, it was surgically removed under anaesthesia (Fig. 11).

**FIGURE 11:**
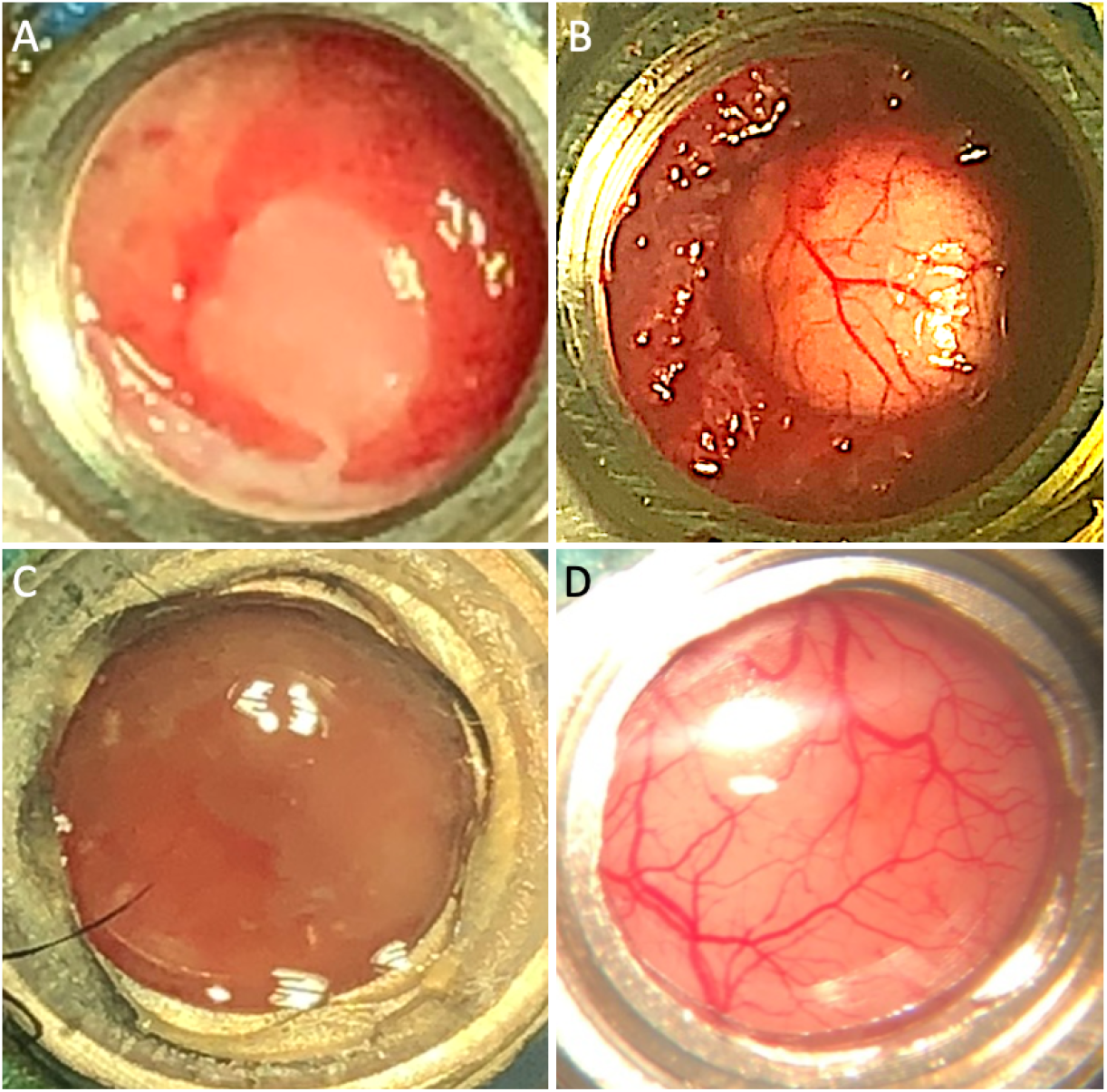
Top: photos of the chamber of the same marmoset as in Fig. 10, before (A) and after (B) resection of the neomembrane. Note the thick “ring” of tissue pushing onto the brain. It had grown starting from the original dura mater that had been left intact in the periphery of the opening. In another monkey (bottom), where the dura had been resected much more extensively, the same phenomenon did not occur, such that, after resection of the neomembrane visible in (C), the cortical surface retained its normal curvature and was completely clear (D). Note that, although images (A,B) have been taken without well, and (C,D) with well (and coverglass), the scale is approximately the same.

In one of the first animals we had removed the dura mater only in the central part of the chamber to an extent matching the well’s central hole, and left it on on the borders of the chamber, the rational being to minimize the risk of haemorrhages due to accidental resection of dural vessels next to the bone. In that particular animal, when it became necessary to remove the neomembrane, we discovered that a thick rim of granular tissue had formed where the dura mater had been left intact. This rim exerted pressure onto the brain increasing its curvature, which, although not causing any detectable symptoms to the animal, severely hampered imaging (Fig. 11A,B). In the subsequent monkeys (Fig 11C,D), we thus removed as much dura mater as possible (essentially up to the edge of the craniotomy), which allowed avoiding this problem.

Following the experience reported in mice (Pisano et al., 2022; Roome et al., 2014; Lefevre et al., 2022), rats (Fenrich et al., 2013), in macaques (Spitler et al., 2008; Namima & al., 2024) and marmosets (MacDougall & al., 2016; Gao & al., 2020), in one marmoset we tried to hinder tissue regrowth immediately after a first surgical neomembrane removal by applying the previously mentioned silicon elastomer “Kwik-Sil” onto the cortex. Its correct application required some dexterity, because care had to be taken to completely dry the cortex before application, and, more difficult, to avoid the formation of bubbles above the region of interest, which seemed to form within the bulk itself of the silicon as it was curing. After mastering these difficulties, a satisfactory visibility could be obtained in the chamber, and Ca-activity could again be imaged in individual cells. Nevertheless, transparency was not as good as in the case where the coverglass was directly touching the cortex. With respect to tissue regrowth, our experience was altogether not conclusive, the silicon seeming neither to accelerate nor to slow down regrowth, although the result might have been different if the silicon had been applied immediately after the first durectomy, before any inflammatory process had started.

Last, when imaging in 2-photon mode, we noticed a slight reddening of the tissue under the silicon elastomer, the origin of which has not been completely elucidated. One possibility could be thermal hyperemia due to the laser beam used for imaging and reduced dissipation of heat due to the isolating properties of the silicon and the absence of cooling by the CSF which is otherwise in contact with the cortex. Additional tests in more animals are needed to clarify the effectiveness of the silicon elastomer against tissue regrowth and its usability in the particular context of two-photon microscopy.

## 4. Discussion

### 4.1 Comparison with previous chamber designs

The need for a cranial recording chamber providing mechanical stability on the one hand, and, on the other, a regular optical interface has appeared already in the early days of optical imaging in the mammalian cortex in-vivo (Blasdel and Salama, 1986; Grinvald et al., 1986). There, the possibility to use long working distance macro-lenses (Ratzlaff and Grinvald, 1991) has given rise to a first generation of imaging chambers that were relatively high (distance between cortex and coverglass in the order of 1-1.5cm), and, being used mostly in macaques (Chen et al., 2002, Arieli et al, 2002) or in acute experiments in smaller animals, relatively large (16-20 mm open aperture) and bulky. Such sizes are problematic for chronic use in smaller animal models such as marmosets, calling for smaller chambers in order for the implantation to be well tolerated. Moreover, with the advent of two photon microscopy *in-vivo*, the chamber’s height had to be considerably reduced in order to adapt to the shorter working distance of typical microscope objectives (at best in the order of a few millimetres). Finally, improved surgical and post-operatory procedures making it possible to image from the same animal for longer and longer times (over several months), underscores the importance of avoiding - or at least slowing down - inflammatory processes and tissue regrowth on the exposed cortex. Experience has shown that neomembrane formation on the cortex can be slowed down by gentle but constant contact with an inert surface, such as glass.

Two recent developments, one by Pattadkal et al, (2025) and the present one, meet this need by using a threaded metal cylinder (“well”) with a coverglass glued to its bottom. By turning it, the height of its bottom glass surface can be easily adjusted to the desired level (although to a different extent in the two designs, see below). Both designs use an outer metal cylinder penetrating into the craniotomy for the whole thickness of the bone, having an external diameter tightly matching that of the craniotomy. Inserting this cylinder into the craniotomy has two advantages. First, it provides additional mechanical stability with respect to previous designs where the chamber was simply “sitting” on top of the craniotomy. Second, together with a resection of the dura mater up to the very border of the craniotomy, this design avoids (or at least slows down) the growth of inflammatory tissue at the periphery of the craniotomy (Fig. 11).

A first difference between the two designs is the use of a curved rather than flat coverglass at the bottom of the well, which warrants better adherence to the cortex than a flat one. Second, the two designs differ with respect to the available field of view. Our chamber provides a clear aperture of 8 mm diameter, twice that provided by Pattadkal’s and colleagues’ design. The latter could obviously be upscaled, yet only to some extent, since 3 mm are added to the chamber’s external diameter due to the bolt needed to fix it to the bone. Height-wise, Pattadkal’s and colleagues’ chamber is only 1.6 mm high, as compared to 3.3 mm in our case. Whereas this has obvious advantages for imaging when using short working distance objectives, accurate machining of such small parts (especially their thread) may be challenging for standard-equipped mechanical workshops as are typically found in research laboratories. Moreover, the chamber’s small height drastically limits the range over which the depth of the glass window can be adjusted in order for it to touch the underlying cortex, also considering that half a millimetre of this 1.6 mm range is lost due to occupation by the protective cap.

Finally, Pattadkal and colleagues endow the chamber’s external cylinder at its very bottom with two winglets extending sidewards below the skull, penetrating between the bone and the dura mater, providing the counterpart to a nut screwed onto the chamber from the (threaded) outside. Whereas this design provides mechanical stability in addition to the dental cement, by pulling from above the bone against the winglets positioned below it, the latter can potentially irritate the dura mater (which at that location is still intact), stimulating inflammation and tissue proliferation. Moreover, due to geometrical constraints, difficulties can arise in positioning the chamber - particularly at locations where the skull’s curvature is asymmetric, potentially resulting in mechanical strain and consequent bone fragility, the marmoset’s skull being only 0.5 - 0.8 mm thick. We thus preferred to design our chamber such that nothing would protrude between bone and dura mater: the part of the chamber entering the craniotomy is smooth and strictly cylindrical, thus minimizing friction against the dura mater. As in Pattadkal’s design, the chamber is finally stabilized using (a small quantity of) dental cement. Until now, none of the implanted animals lost its chamber, nor did any chamber become untight.

### 4.2 Possible improvements

Our design could relatively easily be improved with a number of quite straightforward modifications. An obvious example is to match the curvature of the glass to the specific individual and patch of cortex it is going to be in contact with. It might also be preferable to adjust the external diameter of the chamber to a commercially available trephine saw size. Indeed, such diamond-particle saws cut the bone but not soft tissue, thus reducing the likelihood of damaging the dura mater or even the cortex during the craniotomy.

Somewhat less straightforward yet not less important is the question of how best to warrant a tight and reliable seal between the external cylinder and the well. Adequately meeting this need represents a challenge for our kind of chamber design because some degree of clearance is nearly unavoidable between parts that move one against the other. We tackled this challenge by purposely keeping the fit between the outer cylinder and the well relatively loose in order to be able to create a semi-permanent seal via the injection of Kwik-Sil into the threaded gap between the two parts. If, with some practice, this targeted injection becomes relatively straightforward, some manual dexterity is still required to deal with the tendency of CSF to invade the thread from the inside while Kwik-Sil is being applied from the outside, which can hamper an adequate adhesion of the product to the metal. Furthermore, the thread’s large clearance hampers the precise adjustment of the well’s height, especially in cases where the brain pulsates strongly due to the heartbeat.

One way to address these issues could be to use some slightly elastic material for the outer cylinder or the well, such as hard rubber, Delrin, or the like, and to create a snug fit between the two parts. Moreover, the pitch of the thread between the well and the outer cylinder could be reduced, although this might not be easily achievable with standard equipment: in our case we simply choose the smallest pitch allowed by the equipment we had available, that is, 0.5 mm/turn. Together, these modifications would not only provide a tight seal, but also increase the mechanical stability of the ensemble, the backside obviously being that it would make the thread also more fragile, demanding for particular care when manipulating the two parts.

## Technical Details

### JCaMP injections

We injected either 0.5µL or 1µL per injection site, at an injection rate of 1nL/s. More recently we increased it to 4-10nL/s. The titres we used were the following:

AAV9-mThy1-tTA: 5.66 × 10^12^, mixed in equal parts with AAV9-Tre3-GCaMP7f: 7.93 × 10^12^, or with

AAV9-hSyn-Ribo-jGCaMP8m: 5.5 × 10^12^.

### Jedi-2p injections

Physical parameters of the Jedi-2p injections (volume and rate) were the same as for GCaMP. The plasmide used to produce the JEDI-2p virus was: pAAV-hSyn-JEDI-2P-WPRE. It had been ordered from Addgene, #179464. The viral vector had been produced on-site by the NeuroVir facility: AAV2/9-hSyn-JEDI-2P-WPRE at a titre of 2 × 10^13^ particles/µL. The JEDI-2P sequence is available from GenBank (GenBank: OL542830).

## Supporting information

technical drawings of the chamber

technical drawings of the headpost

technicchamber and headpost, as well as their position on the skull

## Acknowledgments

The authors would like to thank the INT facility Neuro-VIr and Florence Jaouen-Gascon-Gonzalo and Eduardo Gascon-Gonzalo for the preparation of the viral constructs, and Salvatore Giancani and Octavio Ruiz for help during the imaging sessions. This work was funded by the grant ANR-18-CE37-0017-01 “Marmobrain”, the 2020 Equipex+ grant “CIRCUITPHOTONICS”, an INT “fonds d’Investissement” grant, and recurrent funding from the CNRS and Aix-Marseille Université.

## Credits

JM, FC, LR, XD, SR, AL and IV conceptualized the project. IV analyzed the data, supervised and administered the project, and wrote the paper with the help of JM and LR. AL provided solutions for methodological difficulties linked to two-photon imaging in the marmoset. LR, CM and MC developed and implemented the chirurgical methodology. JM and, later, CM performed the microinjections. XD found and implemented the solutions to the mechanical challenges posed by the project, in tight interaction with JM and AL. FC and IV organized the funding for the project.

## Supplementary Material

The technical drawings of the chamber, the technical drawings of the headpost, as well as the technical drawings of their position on a skull are provided as PDFs in the Supplementary Material.

## Highlights

- Cranial chamber for chronic optical recordings in the cortex of small primates
- Individually tailored via MRI guidance and suitable for multiphoton microscopy
- Design enables repeated easy and rapid cortical access and resealing
- Curved glass window with adjustable height delays neomembrane regrowth
- Surgical procedures for chamber implantation are described

