## Supplementary material for "Chamber Implant for Chronic Optical Recordings from the Cerebral Cortex of Marmosets": technical drawings of the chamber

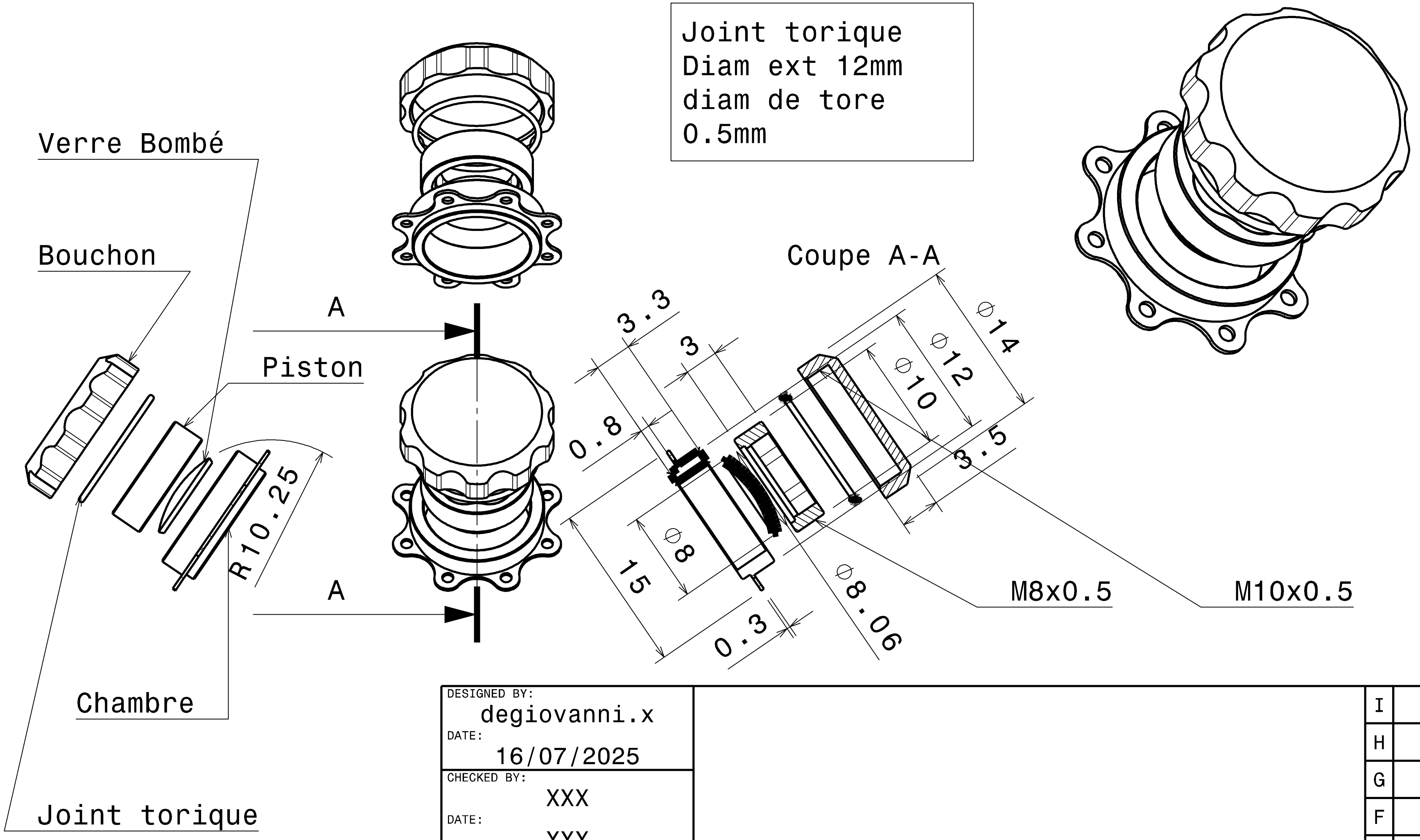


| | |
|---|---|
| DESIGNED BY: | degiovanni.x |
| DATE: | 16/07/2025 |
| CHECKED BY: | XXX |
| DATE: | XXX |
| SIZE | A4 |
| TITLE | Chambre Marmouset |
| SCALE | 1:1 |
| WEIGHT (kg) | 0,01 |
| DRAWING NUMBER | Assemblage chambre marmouset |
| SHEET | 1/1 |

| Rev | |
|---|---|
| I | _ |
| H | _ |
| G | _ |
| F | _ |
| E | _ |
| D | _ |
| C | _ |
| B | _ |
| A | _ |

This drawing is our property; it can't be reproduced or communicated without our written agreement.
