## Supplementary material for "Chamber Implant for Chronic Optical Recordings from the Cerebral Cortex of Marmosets": technical drawings of the headpost

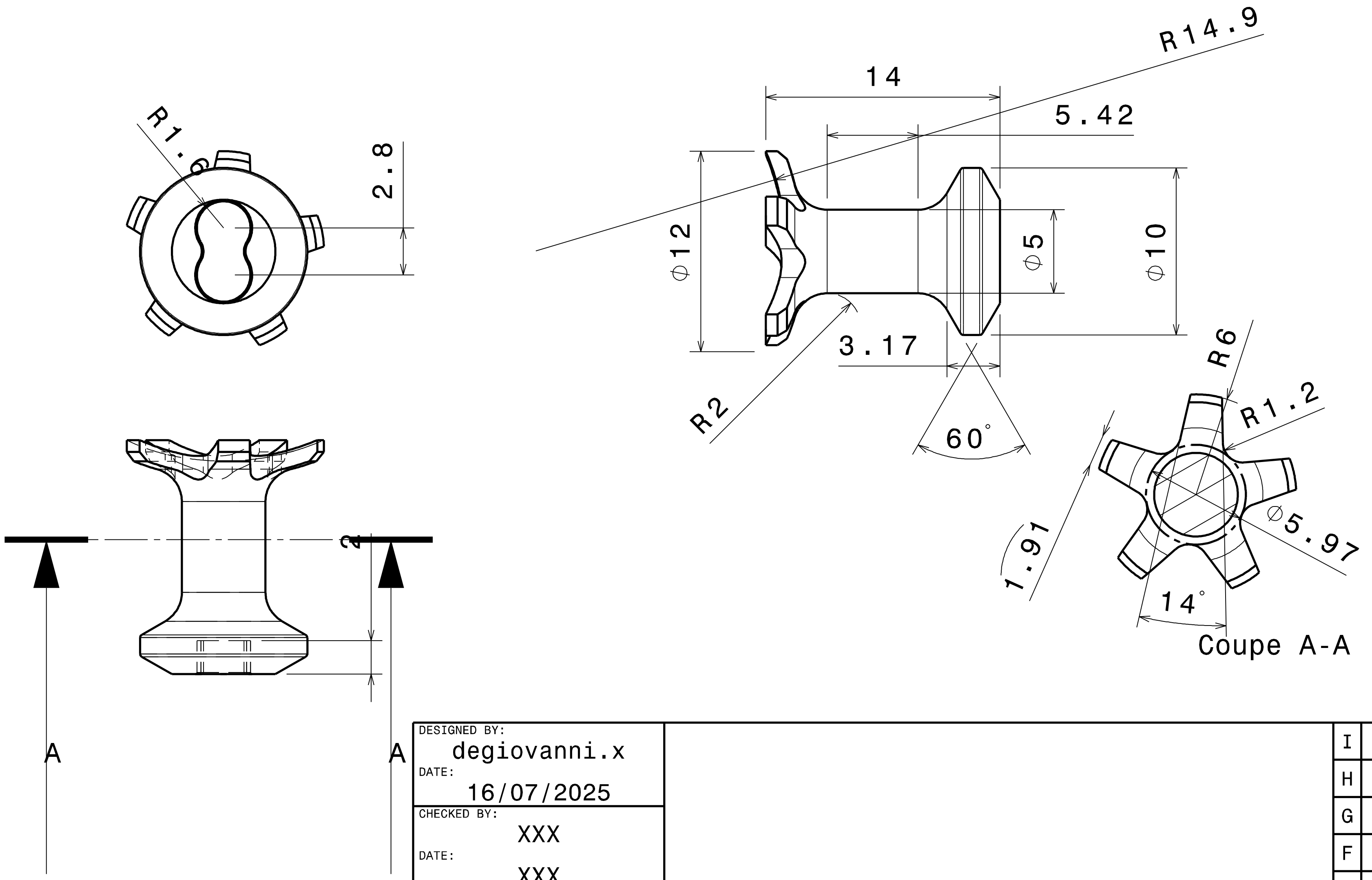


| | | | | | |
|---|---|---|---|---|---|
| DESIGNED BY: degiovanni.x | | | | I | _ |
| DATE: 16/07/2025 | | | | H | _ |
| CHECKED BY: XXX | | | | G | _ |
| DATE: XXX | | | | F | _ |
| SIZE A4 | | Plot Titane Marmouset | | E | _ |
| | | | | D | _ |
| SCALE 1:1 | WEIGHT (kg) 0,00 | DRAWING NUMBER Plot Marmouset | SHEET 1/1 | C | _ |
| | | | | B | _ |
| | | | | A | _ |

This drawing is our property; it can't be reproduced or communicated without our written agreement.
