## Supplementary figures and images for "Chamber Implant for Chronic Optical Recordings from the Cerebral Cortex of Marmosets"

### technicchamber and headpost, as well as their position on the skull

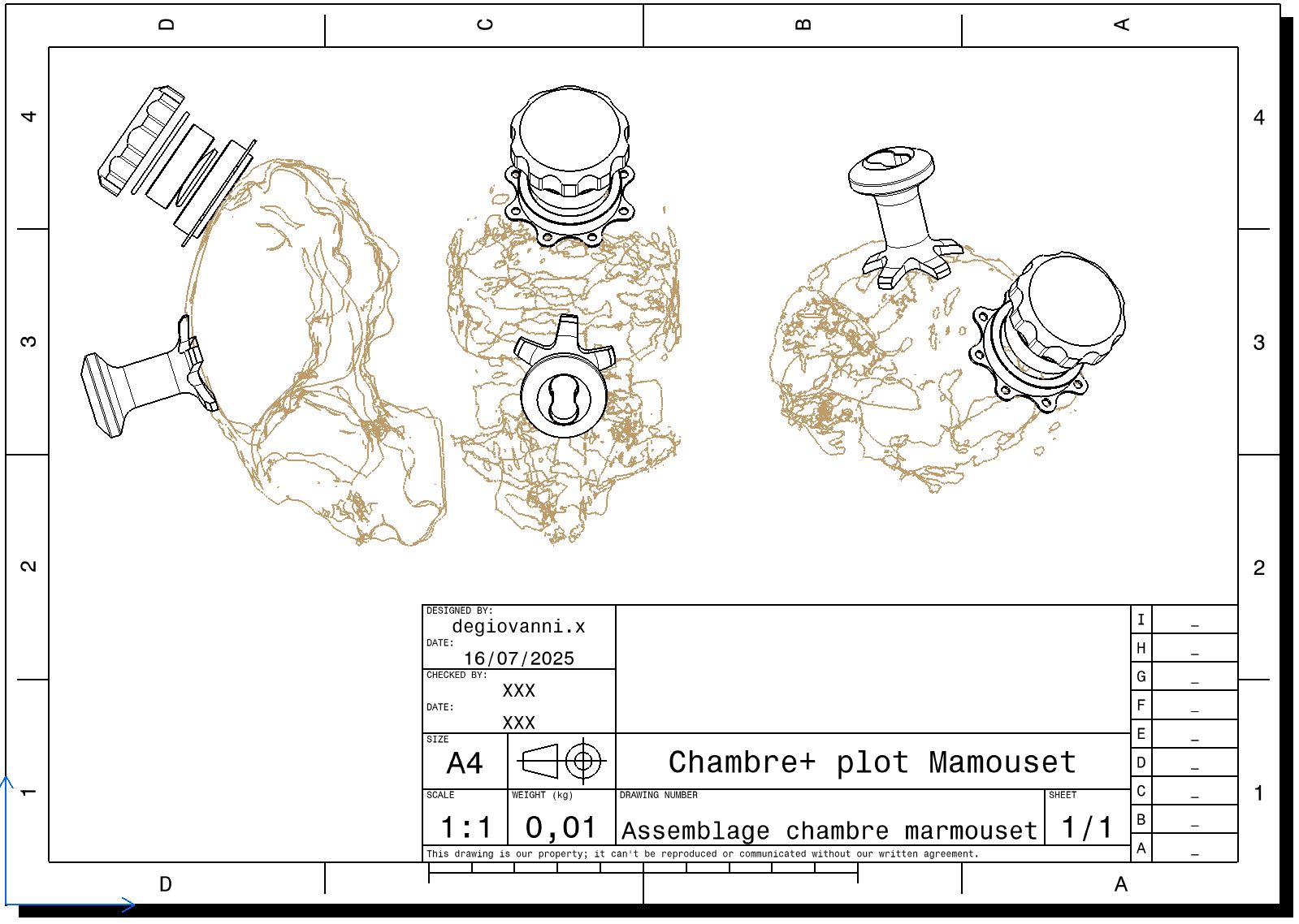
